# Diabetes mellitus and tuberculosis comorbidity induce unique microbial gut dysbiosis and associated metabolome

**DOI:** 10.1101/2025.03.09.642224

**Authors:** Suchitra Jena, Nikhil Bhalla, Shweta Chaudhary, H. Nanaocha Sharma, Pradipta Jana, Bhabatosh Das, Ranjan Kumar Nanda

## Abstract

**Background:** Patients with diabetes and tuberculosis (DMTB) often exhibit atypical disease progression and poor treatment outcomes. Alterations in gut microbiota may contribute to the impaired immune response in diabetes. Understanding the microbiota composition and associated metabolite profiles in DMTB could provide insights for developing targeted adjunctive therapies to improve patient prognosis.

**Method:** Nicotinamide (NA) and streptozotocin (STZ) induced diabetic C57BL/6 mice were aerosol infected with 100-120 CFU of *Mycobacterium tuberculosis* H37Rv. At different time points (2 days pre-infection and 3- and 8 weeks post-infection; wpi), faecal samples were collected for microbiome and metabolome profiling using 16S rRNA sequencing and gas chromatography-time of flight-mass spectrometry (GC-TOF-MS), respectively.

**Results:** Diabetic mice showed gut dysbiosis, with 16 bacterial genera significantly altered between DMTB and TB groups at 3 wpi, which reduced to 11 at 8 wpi. Among the 93 identified faecal metabolites, 11 and 10 metabolites were significantly deregulated (log_2_FC>±1.0, p<0.05) in the DMTB group compared to TB group at 3 and 8 wpi, respectively. A negative correlation between faecal valeric acid levels in DMTB with the abundances of *Coriobacteriaceae, Granulicatella, Veillonella, Achromobacter, Erysipelatrichaceae, Atopobiaceae, Citrobacter*, and *Lactiplantibacillus* were observed. Stronger correlations between microbiota and metabolite were observed in DMTB comorbidity compared to TB. Additionally, correlation analysis also revealed a phylogenetic distance-dependent synchrony in amino acid metabolism across five clades, each containing three or more bacterial genera.

**Conclusion:** This study reveals significant differences in the gut microbiome and metabolome between DMTB and TB groups. The identified microbes and metabolic products may serve as potential prebiotic or probiotic candidates for improving recovery in DMTB comorbid conditions.

## Introduction

Diabetes mellitus (DM) is a metabolic disorder characterised by high blood glucose levels, showed increased case incidence from 200 million in 1990 to 830 million in 2022 (1, 2). Patients with DM exhibit increased susceptibility to infectious diseases such as tuberculosis (TB), often presenting with more severe symptoms that are difficult to manage clinically. *Mycobacterium tuberculosis* (Mtb), the pathogen responsible for TB, shows faster growth in the presence of sugar alcohols like glycerol in vitro. Diabetic patients with higher glycerol abundance probably facilitate faster Mtb growth, leading to a more severe form of TB (3). Due to different metabolic phenotypes, the diabetes and tuberculosis (DMTB) and TB cases may present different disease pathologies, and detailed studies are missing.

Metabolic changes associated with diabetes lead to low-level inflammation, increased oxidative stress, and impaired immune cell function and infiltration (4). DMTB subjects have relatively higher impairment in cell-mediated immunity and cytokine response, neutrophil inflammation, and alveolar macrophage responses. These factors contribute to higher morbidity in DMTB subjects, relapse rate and drug resistance development (5). The impact of DM on the digestive system causes delayed gastric emptying in the upper gastrointestinal system and the small intestine, diabetes-associated enteropathy through vagal nerve dysfunction and a negative impact on Cajal and nNOS interstitial cells (6–8). The slowed peristalsis in DM is often the cause of already-known small intestinal bacterial overgrowth, which can potentially affect the gut microbiota (6–8)

Gut microbiota plays a crucial role in shaping immune development in infants and maintaining host health in adults by modulating metabolism and immune responses. A healthy gut microbiota comprises six phyla: *Firmicutes, Bacteroidetes, Actinobacteria, Proteobacteria, Fusobacteria*, and *Verrucomicrobia* (10). Gut dysbiosis associated with DM leads to the overgrowth of opportunistic pathogens belonging to *Bacteroides, Clostridium, Eggerthella*, and *Escherichia* and is linked with impaired beta cell function, insulin sensitivity, and glucose metabolism (9–11). A decreased abundance of *Clostridiales* and a higher abundance of *Pophyromonodaceae* with treatment were reported in TB patients (12,13). Investigating the influence of diabetes on gut microbiota and associated metabolite profiles in subjects infected with Mtb could provide insights into the disease etiology and develop microbe or metabolites-based concoctions for better disease management.

This study compares faecal microbiota and metabolite profiles of DMTB and TB groups at various time points. These findings offer valuable insights for developing prebiotic and probiotic formulations of bacterial cultures and metabolites to improve management of DMTB comorbid conditions.

### Methodology

### Diabetes induction and Faecal matter collection

The animal experimental protocols in this study were approved by the animal ethics committee (vide reference ICGEB/IAEC/07032020/TH-13Ext) of the International Centre for Genetic Engineering and Biotechnology New Delhi component. Faecal samples were collected from the mice used for another study (14). Briefly, male C57BL/6 mice aged 8-12 weeks were administered nicotinamide (60 mg/kg in 0.9% saline; NA) and 15 minutes later streptozotocin (150 mg/kg in 50 mM citric acid buffer; STZ) 3 times at an interval of 10 days each. To establish DM, blood glucose levels were monitored for several weeks after the last NA-STZ dose (14). Fresh faecal pellets (at least 2-3 per mouse) were collected in containers directly from the mice after gentle pressing and stored at -80°C until further processing.

### Faecal metabolite extraction and derivatisation for GC-TOF-MS data acquisition and data processing

To the faecal samples (50 mg), zirconia beads (300 mL, 0.1 mm diameter), chilled methanol (80%, 1 mL) and ribitol (0.5 mg/mL, 2 mL) were added and subjected to bead-beating (3 cycles) using BioSpec Bead beater. The homogenised faecal sample was centrifuged at 16,000 g at 4 °C for 15 minutes, and the supernatant was harvested. A fraction of the extracted faecal metabolite (400 mL) was vacuum-dried using SpeedVac (Labconco, USA) at 40 °C for complete dryness to treat with methoxamine–hydrochloric acid mix (40 μL, MOX-HCl, 20 mg/mL) for 2hrs at 60 °C, 900 rpm. After that, the reaction mixture was incubated with N-methyl-N-(trimethylsilyl) trifluroacetamide (MSTFA, 80 mL) and trimethylchlorosilane (TMCS;1%) for 30 min at 60 °C at 900 rpm. After centrifugation at 16,000 g for 10 min at room temperature, the supernatant was harvested and transferred to GC vials for GC-TOF-MS acquisition.

### GC-TOF-MS and data processing

Using an automated multipurpose sample introduction system, the derivatised samples were loaded into the column attached to the GC-TOF-MS instrument (Agilent Technologies, Santa Clara, CA) in split-less injection mode. Helium was used as a carrier gas with a 0.5 ml/min flow rate for each run. Metabolite separation was carried out in an RTX-5 column (30 m, 0.25 mm, 0.25 mm) with a temperature gradient from 60-300°C at a ramp rate of 10°C/min (60-220°C) and 5°C/min (220 to 300°C). Hold times of 1 min and 5 min were kept at the start and end of the run, respectively. The injection port temperature was held at 250°C throughout the run. Mass spectrometric data acquisition was carried out at -70 eV, and a mass range of 50-550 was scanned with a rate of 50 spectra per second. A 600-sec solvent delay was used as the GC-MS parameters and was controlled using ChromaTOF software (version 4.50.8.0; Leco, USA). Raw GC-MS data files (n=103), including all QC and study samples, were processed and aligned using the “Statistical Compare” feature of ChromaTOF. The minimum peak width was set at 1.3 sec for peak picking, and the signal-to-noise ratio (S/N) threshold was 75 for tentative molecular feature identification, mainlib, nist_ri, nist_msms2, nist_msms, and replib libraries from NIST were used. The maximum retention time difference was set at 0.5 sec, and the minimum spectral similarity was set at 600 for the spectral match. Aligned peak information was exported in “.csv” format, and molecules absent in more than 50% of samples of at least one class were excluded from the analysis. Before statistical analysis, manual data curation was carried out to align peaks and remove unsilylated molecules and silanes from the data matrix.

### Total DNA extraction from faecal samples

The faecal sample (150-200 mg) were subjected to homogenisation through vortexing in 250 μL of Tris_50mM_EDTA_1mM_ buffer (pH 8.0) with 4 glass beads (2.7mm). The homogenates were treated with 50 μL of lysozyme (10 mg/mL), 50 μL of mutanolysin (3KU/mL) and 3 μL of lysostaphin (4KU/mL) for 1 hour at 37°C. 300 μL of 4 M guanidine thiocyanate (GITC) were added and mixed for 45 seconds. Next, 300 μL of 10 % N-lauryl sarcosine were added, incubated for 10 mins (300 rpm, 37°C), mixed and incubated (1hr, 70°C) followed by two cycles of bead beating (with 300mg of zirconia, 0.1 mm). Poly(vinylpolypyrrolidone) (PVPP, 15mg) was added and mixed, followed by centrifugation at (16,000 g, 3 mins). The supernatant was saved, and the pellet was washed twice with 500 μL of Tris(50mM)-EDTA(20mM)-NaCl(100mM)-PVPP(1%) buffer, centrifuged, and the supernatant was pooled. The pooled supernatant was centrifuged (16,000g, 3 mins). To the resulting supernatant,4 mL of isopropanol was added and incubated at room temperature for 10 mins, followed by centrifugation (16,000g, 10 mins). The supernatant was discarded, and the pellet was dried for 1hr at 42°C. To the dried pellet, phosphate buffer and potassium acetate (1.0 mL, 9:1) were added and incubated overnight at -20°C. The samples were centrifuged (16,000g, 30 minutes, 4°C). RNase A (4 μL, 10 mg/mL) was added to the supernatant and incubated for 30 minutes at 37°C. To the supernatant, 100 μL of 3M sodium acetate was added, followed by the addition of ice-cold ethanol (96%), mixed, and subsequently incubated for 5 minutes at room temperature. The samples were centrifuged (16,000g, 15 minutes) at 4°C, and the pellet was washed with ice-cold ethanol (70%). The pellets were dried in an incubator for 1hr at 42°C. The dried pellets were then resuspended in 150 μL of nuclease-free water (NFW) and incubated for 10-15 mins at room temperature, followed by centrifugation at 16,000g for 10 mins at 4°C. The supernatant was stored and quantified using a Qubit 3.0 Fluorometer.

### Enrichment of 16S rRNA and Sequencing

V3 and V4 regions of the 16S rRNA gene were amplified using the following primers: Forward Primer: *5’-TCGTCGGCAGCGTCAGATGTGTATAAGAGACAGCCTACGGGNGGCWGCAG-3* and Reverse primer: *5’ GTCTCGTGGGCTCGGAGATGTGTATAAGAGACAGGACTAC HVGGGTATCTAATCC-3’*. The amplification was done using KAPA high-fidelity polymerase. PCR products were purified and quantified using the Qubit dsDNA quantitation method (Invitrogen Inc, USA). For the preparation of the NGS library, the amplified DNA was subjected to Tagmentation using a NexteraXT kit (Illumina Inc, USA). Sequencing was carried out in the 300×2 paired-end configuration on the Illumina NextSeq2000 platform (Illumina, USA).

### Data processing

The sequencing reads were subjected to rigorous processing using NF-core/ampliseq pipeline v2.12.0v. NF-core/ampliseq pipeline included DADA2-based ASV calling, and a taxonomy assignment was carried out. SILVA 138 database was used to annotate the amplicon sequence variants. The ASV matrix and taxonomy files were subsequently analysed using QIIME2 v2024.10.1. The analysis was done at the genus level. The ASVs that were unclassified at the genus level were summed up and studied at the family level.

### Clustering of genera and molecules

For genera, the 16S rRNA sequences were extracted from the taxonomy file from DADA2 output and aligned with clustal omega and the Newick tree was calculated with FastTree. For metabolites, the SMILES were used to generate fingerprints, which were used to determine the distance between molecules. The Newick tree generated from these was used to cluster correlation matrices between metabolite and microbiota.

### Statistics

The metabolite data were normalised to the peak areas of the internal standard (ribitol). A pseudocount of 1 was added before the log_2_ transformation, and the log-transformed matrices were used for PCA analysis. The paired student’s t-test was performed for differential abundance, and metabolites that showed log_2_FC > ± 1 with raw p< 0.05 were considered statistically significant. For microbiome analysis, pairwise comparisons were done using the Deseq2 package, and ANOVA and Kruskal-Wallis tests were used for multi-group analysis. A pseudocount of 1 was added to Deseq2 normalised counts before log transformation of the microbiome ASV matrix. The signals were converted to Z-scores, and pairwise Pearson’s correlation coefficient was computed. Fisher’s Z-test was carried out using Python libraries: numpy, pandas, scipy.stats.norm for differential c correlation analysis. The results were parsed in MS-Excel 2016 home edition. The figures were created using Python and GraphPad Prism 8.4.

## Results

### Diabetes induction and experimental design to determine the effects on faecal metabolome and microbiome in Mtb-infected mice

The mice used in this study were the same set used and described previously, with the detailed diabetes validation metrics (14). Briefly, a set of C57BL/6 male mice (8-12 weeks) received a nicotinamide-streptozotocin (NA-STZ) combination and developed hyperglycemia and dyslipidemia before aerosol infection with low dose (100-120 CFU) of Mtb H37Rv using a Madison Aerosol Chamber in BSL-III aerosol challenge facility at the host institute (Figure 1).

**Figure 1:**
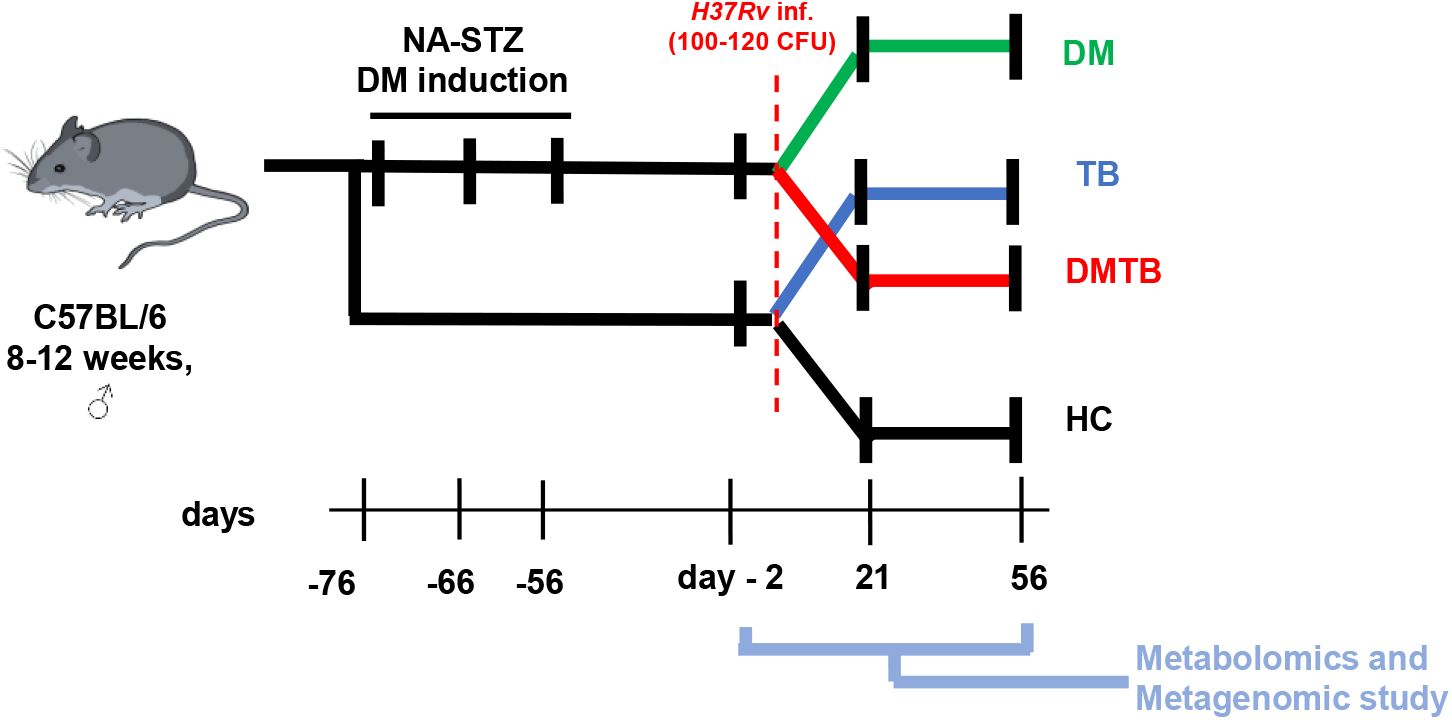
Experimental plan adopted in this study. Diabetic induction in C57BL/6 male mice (8-12 weeks old) was done using the nicotinamide-streptozotocin (NA-STZ) combination. Diabetes was confirmed with consistently elevated blood glucose levels after the last NA-STZ cocktail dose. A faecal sample was collected 2 days before and 3 and 8 weeks after Mtb infection. The infection was carried out through aerosol mode. Note: The mice used in this study were the same as those described previously, and diabetes validation can be found in our earlier published work (14). DM: diabetic, TB: Mtb infected, DMTB: Diabetic and Mtb infected (comorbid), HC: Healthy controls, NA: Nicotinamide, STZ: Streptozotocin, C57BL/6: mice strain.

### Impact of diabetes on gut microbiome dynamics

To investigate microbiome dynamics in hyperglycemic and euglycemic mice at different time points, 16S rRNA sequencing was performed (Figure 2A). The PCoA plot of euglycemic mice showed a time-dependent change but was minimal in DM mice (Figure 2B). Alpha diversity, as assessed by Shannon’s index, was significantly higher in the diabetic mice compared to the healthy controls at week 3. Additionally, a time-dependent decrease in alpha diversity variance was observed (Figure 2C). To determine the microbes showing significant changes in their abundance, a multi-group analysis using one-tailed ANOVA was performed. The analysis revealed time-specific perturbations between groups (p < 0.05), leading to the selection of 22 microbiota, which were further categorized into five trend-specific clusters. Cluster-I, included *Dubosiella, Akkermansia* and *Erysipelatoclostridium*, which were lower in abundance in DM, while in healthy, their abundance decreased over time (Figure 2D). Cluster-II, included *Parvibacter, Parasutterella, Muribaculum, Family XIII UCG-001, Butyricicoccus* and *Olsenella*, which remained similar in DM, unlike the healthy (Figure 2E). Cluster-III, included *Serratia, Harryflintia, Deslfovibrio* and *Atopobiaceae*, and their abundance was lower in the DM group (Figure 2F). Cluster-IV included *Weissella, Limosilactobacillus, Lachnospiraceae UGC-0* and *Enterobacter*, and their abundance remained low in healthy but present in DM (Figure 2G). All genera in Cluster-V showed a unique pattern. *Alloprevotella* abundance was low in DM at both time points. UCG-010 showed inverse abundance patterns between the DM and the healthy group. DNF0089 showed a time-dependent increase in both the healthy and diabetes groups, with a slower rate in DM. *Odoribacter* showed a decrease in abundance at week 3, followed by a minor increase at week 8 in the DM group. In healthy. *Allobacculum* showed a stable abundance close to zero in the DM group, and a time-dependent increase in trend was observed in healthy (Figure 2H). These findings demonstrate that diabetic mice exhibit significant changes in microbial abundance over time, which could impact their microbiota-associated metabolic profiles.

**Figure 2:**
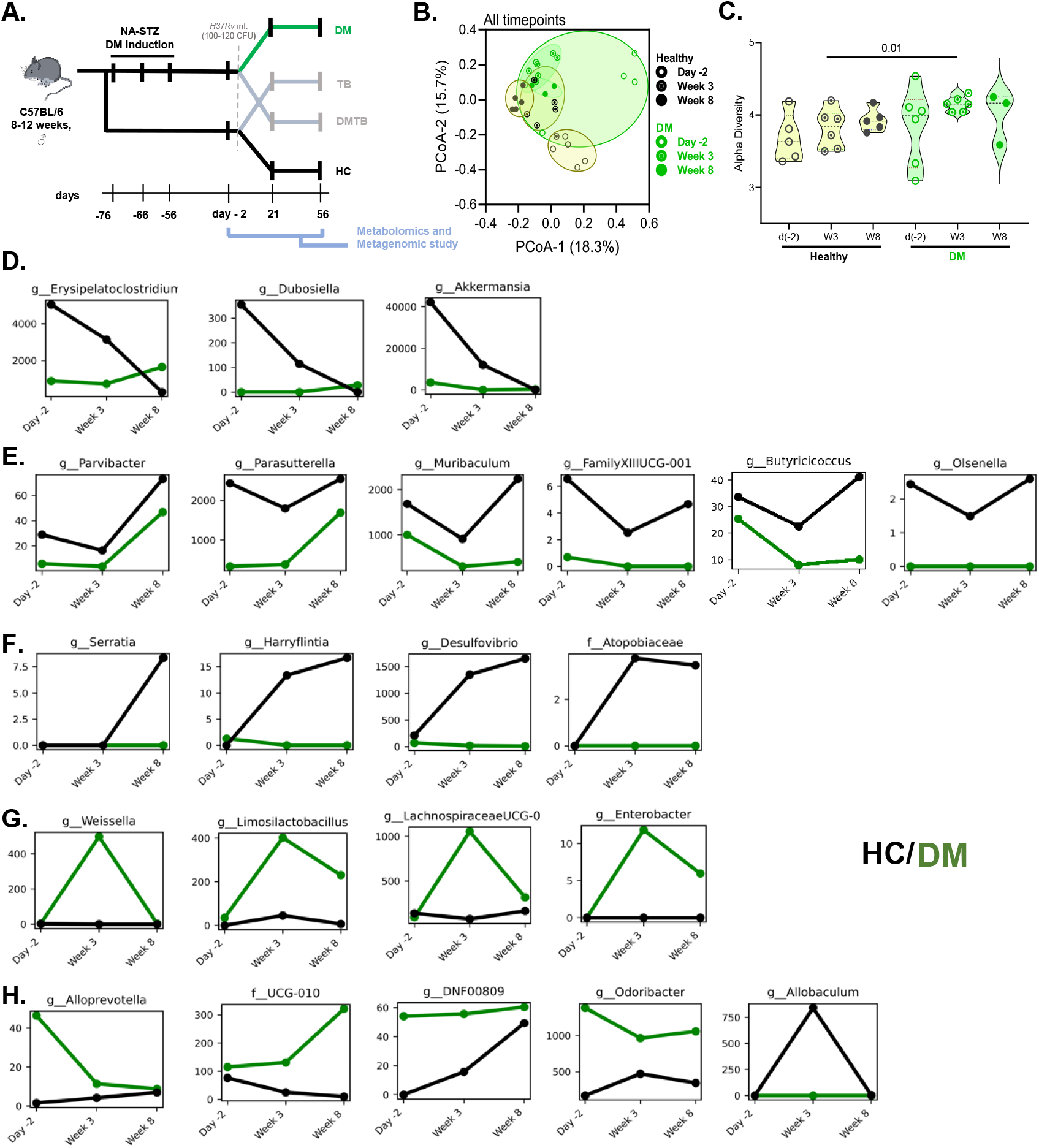
Faecal microbiome changes are impacted by diabetes. Diabetes was induced using an NA-STZ, and faecal matter was collected at three-time points. 16S rRNA sequencing of the faecal sample was performed as mentioned in the methodology, and Deseq2 normalized abundances were used for each genus group-wise and microbiome differences at different time points. Species that showed p<0.05 (One-way ANOVA) were down-selected, and their abundances were averaged for plotting against time points. A: PCA plot. B: Genera showed a decline after Mtb infection in healthy mice. C: miscellaneous trend observed. D: Genera showed a decrease at week 3 and an increase at week 8 after Mtb infection in Healthy with a similar but less pronounced effect in DM mice. E: Bacteria showed an increase only in healthy mice. F: Genera showed a sudden spike in week 3 and a sudden decrease in week 8 after the Mtb infection. G: Miscellaneous trend group 2. g: genus; f: family, DM: diabetic, TB: Mtb infected, DMTB: Diabetic and Mtb infected (comorbid), HC: Healthy controls, NA: Nicotinamide, STZ: Streptozotocin, C57BL/6: mice strain.

### Diabetes-driven changes in faecal metabolites influence outcomes of Mtb infection

To understand DM-associated metabolite differences in Mtb-infected mice, we compared the faecal metabolites between the DMTB and TB groups (Figure 3A). The comparative metabolite differences are presented in the supplementary information (Figure S2). The PCA plot of DMTB and TB groups at 3 and 8 wpi showed partially overlapping clusters (Figure 3B and 3C). At 3 wpi, 11 deregulated (3/8: up/down; log_2_FC > ±1.0, students t-test p < 0.05) metabolites were observed in the DMTB compared to the TB group. Increased abundance of methyl-alpha-D-glucopyranoside, 2-hydroxyadipic acid, 4-methylmandelic acid, 2-hydroxyvaleric acid, L-leucine, L-tryptophan and sugars such as D-(−)-ribofuranose and D-xylopyranose and a lower abundance of pseudo uridine, pipecolinic acid, phosphoric acid was observed (Figure 3D). At 8 wpi, a set of 10 deregulated metabolites was identified, out of which an abundance of L-tryptophan, GABA, 5-amino valeric acid, D-pinitol, 5-hydroxyindole-3-acetic acid and xylulose were higher in DMTB than TB. In contrast, a low abundance of L-leucine, pipecolonic acid, valeric acid and 3-hydroxyphenylpropionic acid was observed in the DMTB group (Figure 3E). At 3 wpi, pseudo uridine was significantly low in DMTB compared to TB, but at 8 wpi, it was lower. Higher L-tryptophan levels were observed in the DMTB group at both time points, indicating its association with diabetes. L-leucine in the DMTB group was higher at 3 wpi, and reduced at later time points, i.e. 8 wpi, compared to the TB group. At 3 wpi., the DMTB group showed higher 2-hydroxy adipic acid (2-HAA) abundance compared to TB and a similar trend was observed between DM and healthy (Figure S1D). Pipecolinic acid, phosphoric acid, pseudo uridine and D-pinitol decreased over time in the TB group. A decreased abundance of L-leucine and higher 5-amino valeric acid and L-tryptophan levels were observed in the DMTB group with time (Figure 3F). These results suggest that DM significantly alters the abundance of specific faecal metabolites, likely driven by changes in both host and microbiota metabolism.

**Figure 3.**
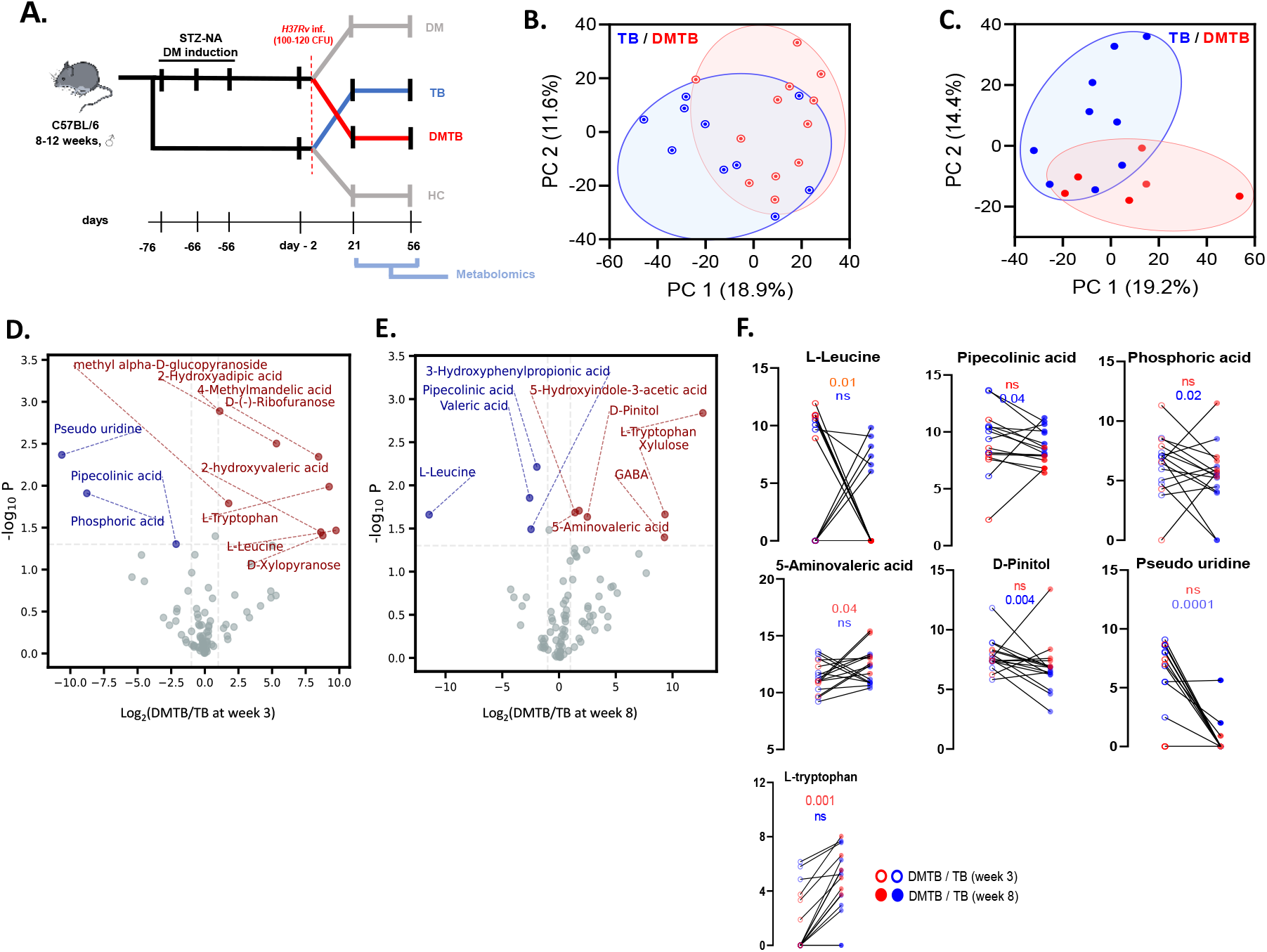
Diabetes-driven changes in faecal metabolites influence outcomes of Mtb infection: The metabolome matrix was normalized to the internal standard and log-transformed before being subjected to PCA and subsequently analyzed. A: Schematic representation of the experimental plan. B; C: PCA plot showing the clustering of the faecal metabolites at week 3 and week 8, respectively. D; E: Differential abundance analysis of metabolites of DMTB vs TB at weeks 3 and 8 post-Mtb infection. F: Levels of deregulated molecules in DMTB vs TB comparisons at weeks 3 and 8 (L-leucine, pipecolinic acid, phosphoric acid, pseudouridine 5-amino valeric acid, D-pinitol, and L-tryptophan). DM: diabetic, TB: Mtb infected, DMTB: Diabetic and Mtb infected (comorbid), HC: Healthy controls, NA: Nicotinamide, STZ: Streptozotocin, C57BL/6: mice strain, PCA: Principal Component Analysis, P: p-value.

### Faecal microbiome of hyperglycemic and euglycemic mice alters upon Mtb infection

To determine the influence of diabetes upon Mtb infection, a comparative microbiome profiling between DMTB and TB was carried out at two-time points, i.e. 3 and 8 wpi. The microbiota of DMTB and TB groups clustered separately, indicating a significant difference in their beta diversity at 3 and 8 wpi (Figure 4A, 4B). The healthy controls were clustered away from the DM group but had similar beta diversity with healthy controls. The alpha diversity of the DM group showed a subtle and significant (p<0.01) increase compared to healthy controls (Figure 4C). Although the DMTB group showed a subtle (p=0.06) increase in alpha diversity from healthy control and was similar with TB. The TB group had lower alpha diversity than the DM group at 3 wpi. At 8 wpi, the differences in alpha diversity were insignificant between all study groups (Figure 4D).

**Figure 4.**
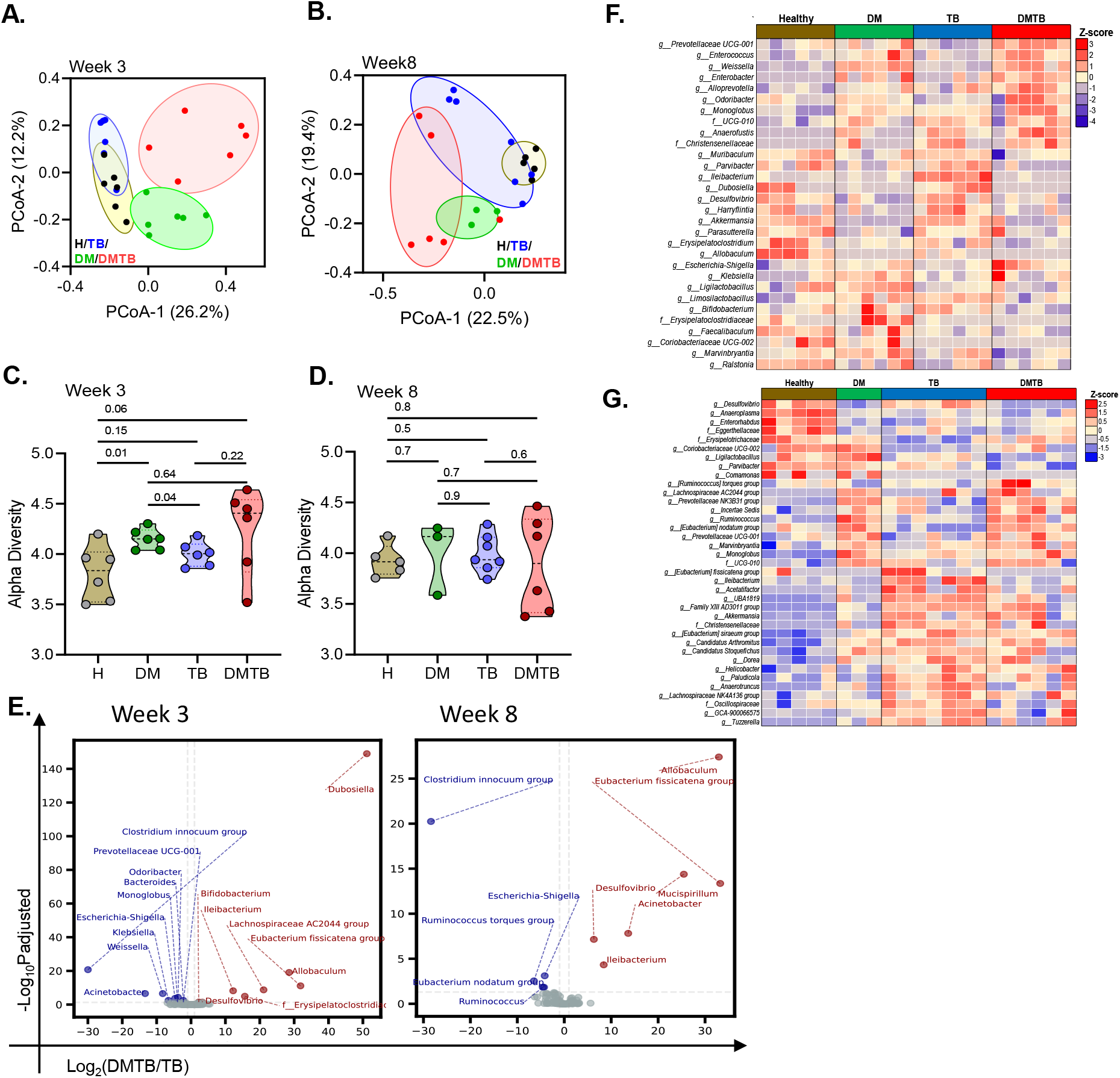
Faecal microbiome changes are impacted by diabetes after Mtb infection. Diabetes was induced in mice using the NA-STZ method and faecal gDNA isolated at 3 and 8 weeks post Mtb aerosol challenge were used for microbiome analysis by using 16S rRNA V3 and V4 specific primers. The sequencing library was prepared using a Tagmentation-based method, and 300×2 PE Illumina chemistry was used for sequencing. The sequencing data was subjected to NF-core/ampliseq standardized pipeline. DADA2 computed ASV matrix was normalized using edgeR for PCA analysis, and normalization was done with Deseq2 to determine microbes undergoing dysbiosis. A: PCoA plot at 3 weeks. B: PCoA plot at 8 weeks (Braycurtis Index). C: Alpha diversity at 3 weeks. D: Alpha diversity at 8 weeks (Shannon). E: Differentially abundant genera between DMTB and DM and TB at week 3 and week 8 post-infection. F and G: The Deseq2 normalized data converted to the first log2-transformed matrix, and species absent in 1/3^rd^ of the samples were excluded from the analysis. Finally, the Kruskal-Wallis test was applied to each species to assess statistical significance across the groups (Healthy, DM, TB and DMTB). Species that showed p_adj_<0.05 have been shown for week 3 and week 8 post Mtb infection. DM: diabetic, TB: Mtb infected, DMTB: Diabetic and Mtb infected (comorbid), HC: Healthy controls, NA: Nicotinamide, STZ: Streptozotocin, C57BL/6: mice strain, PCA: Principal Component Analysis, P: p-value.

To capture time-dependent changes in the microbiome profile of DMTB and TB groups, a pairwise analysis between 3 and 8 wpi was carried out. The abundance of *Allobaculum, Desulfovibrio, Ileibacterium* and *Eubacterium fissicatena group* was higher in the DMTB group. Similarly, an abundance of the *Clostridium innocuum group* and *Eschereichia-shigella* genera were observed to be low in the DMTB group compared to the TB group. Whereas we observed significant variations in bacterial genera, at 3 wpi, abundance of *Clostridium innocuum group, Prevotellaceae UGC-001, Odoribacter, Bacteroides, Monoglobus, Klebsiella, Weissella* and *Acinetobacter* were low and higher *Dubosiella, Bifidobacterium, Lachnospiraceae AC2044*, and also a family i.e. *Erysipelatoclostridiceae* (Figure 4E top right). At 8 wpi, a higher abundance of *Mucispirillum* and a lower abundance of *Ruminococcus* and *Eubacterium* groups was observed in DMTB (Figure 4E Bottom right). A multigroup analysis yielded similar findings observed in the pairwise analysis (Figures 4F and 4G, Supplementary Figure S2). These findings indicate that DM induced severe microbial gut dysbiosis in Mtb infected study group.

### Microbiome-metabolite correlation analysis reveals phylogenetic distance-dependent synchrony in gut microbiota amino acid metabolism

Diabetic condition perturbs both the host microbiome and metabolic profile. We aimed to correlate the gut microbiome and the deregulated metabolome, in the DMTB group and assess how these changes differ from TB. Multiple correlation clusters were observed in the DMTB comorbid group compared to the TB (Figure 5A, B,C,D). L-tryptophan, which was higher in DMTB, showed positive correlations at 3 wpi with *Prevotellaceae, Succinivibrio, Acinetobacter, Faecalibacterium, Neisseria* and at 8 wpi with *Paludicola, Halomonas, Enterococcus, UCG-005, Rikenella*. A higher abundance of L-leucine at 3 wpi was positively correlated with *Anaerococcus, NK4A214 group*, and *Eubacterium coprostanoligenes group*. At 8 wpi, *Anaerococcus* (r=-0.16), *NK4A214 group* (−0.57) and *Eubacterium coprostanoligenes group* (r=-0.45) showed a negative correlation with L-leucine. Pipecolinic acid levels were low in DMTB, were negatively correlated (r=-0.7 or 7.0) with *Prevotellaceae, Roseburia, Escherichia-Shigella, HT002* and at 8 wpi correlated with *Ileibacterium, Anaerofilum, Erysipelotrichaceae* (Figure S3). Differential correlation analysis of microbiome-metabolome of DMTB comorbid and TB showed higher deregulated clusters at 3 wpi. *Atopobiaceae, Achromobacter, Anaerofilum, Citrobacter, Coriobacteriaceae UCG-002, Erysipelotrichaceae UCG-003, Granulicatella, Lactiplantibacillus, Veillonella* showed a negative correlation (r<-0.9, p_adj_<0.001) with valeric acid in DMTB group compared to TB group at 3 wpi. *Christensenellaceae r-7 group* and *Lachnospiraceae AC2044* group negatively correlated (r<-0.9, p_adj_<0.001) with citric acid in DMTB than in the TB group. *Anaerofilum* and *Lactiplantibacillus* negatively correlated (r<-0.9, p_adj_<0.001) with d-(+)-ribono-1,4-lactone. This indicates that the metabolic states of bacteria from different genera are in higher metabolic synchrony in DMTB compared to the TB group (Figure 5B). At 8 wpi, *Porphyromonas*, and *Ruminiclostridium* positively correlated with L-norleucine (r>0.6, p_adj_<0.001), indicating their synchronised metabolic states (Figure 5C). In addition, a subset of amino acids and their derivatives showed correlations in DMTB at 8 wpi (Figure S3D). A filtered heat map of these amino acids showed 5 clusters, each consisting of 3 or more genera, correlated with amino acid levels (Figure 5G). These findings suggest substantial correlations between microbiome and metabolome in DMTB groups and possible synchrony in their metabolic states in a phylogenetic distance-dependent manner in a genera-specific way.

**Figure 5:**
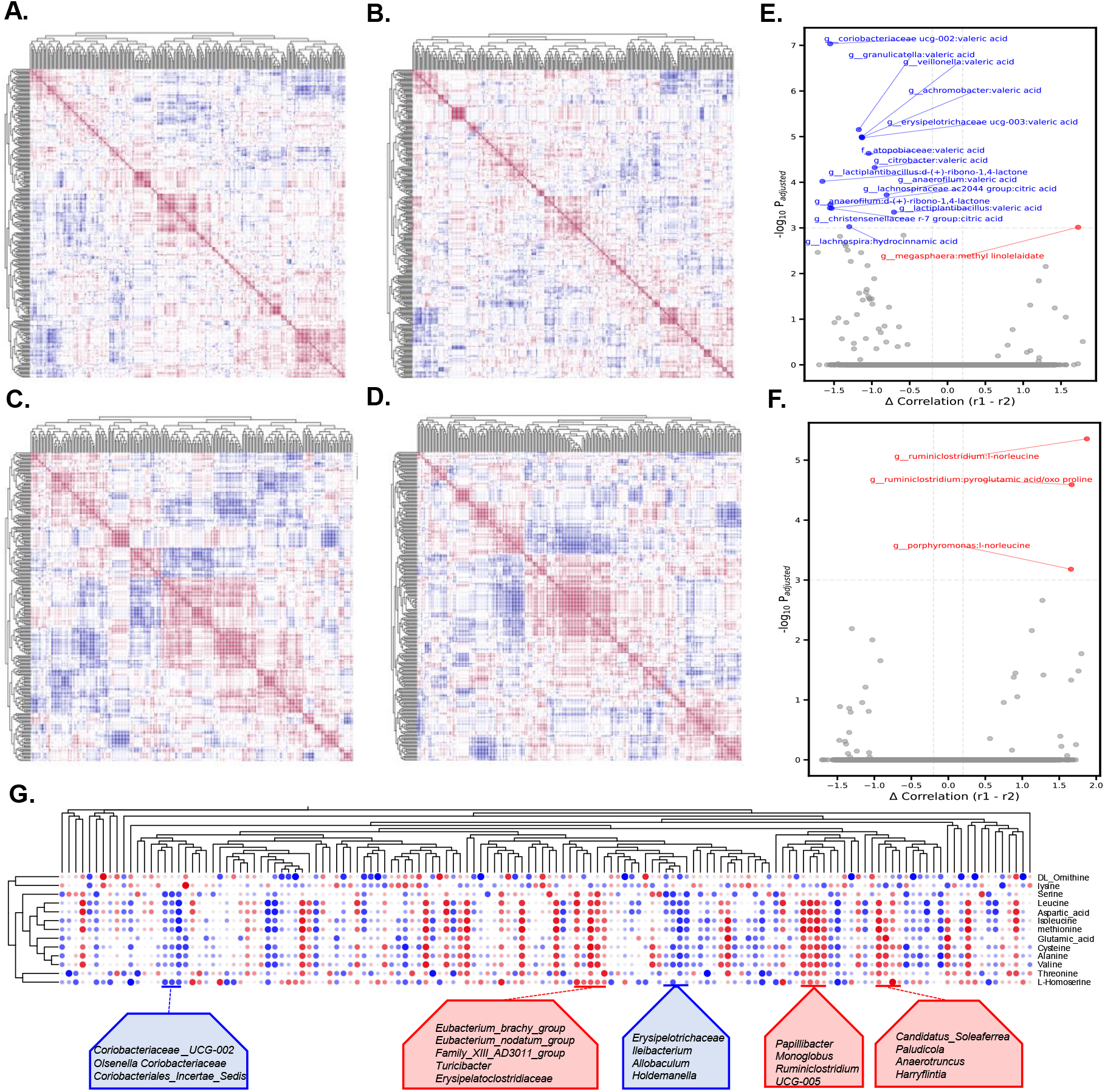
Metabolite-microbiome correlation analysis revealed a higher correlation in the DMTB group and phylo-distance specific metabolic synchrony in amino acid metabolism in some genera. A; B; C; D: Deseq2 normalized abundance and metabolite abundances were log-transformed and converted to Z-scores for correlation analysis within and between the microbiome and metabolome. The pairwise Pearson correlation coefficient between microbiome and metabolome was computed for DMTB and TB in weeks 3 and 8 post-Mtb infection. Correlation between microbes and metabolites is represented in the form of heatmaps at TB at week 3 (A), week 8 (B), DMTB at week 3 (C) and week 8 (D). E; F: Differential correlation analysis using Fisher’s Z-test. The correlation differences with p_adj_ <0.0001 were considered significant. Differential correlation analysis between DMTB and TB at week 3 (E) and week 8 (F). G: Filtered correlation matrix showing clusters of bacteria with similar correlation with amino acids indicating metabolic synchrony. Note: If r1 -r2 > 0, the correlation is more positive in DMTB than in TB, and if r1 -r2 < 0, the correlation is more negative in DMTB than in TB. Abbreviations: r1: correlation coefficient in DMTB; r2: correlation coefficient in TB.

**Figure 6:**
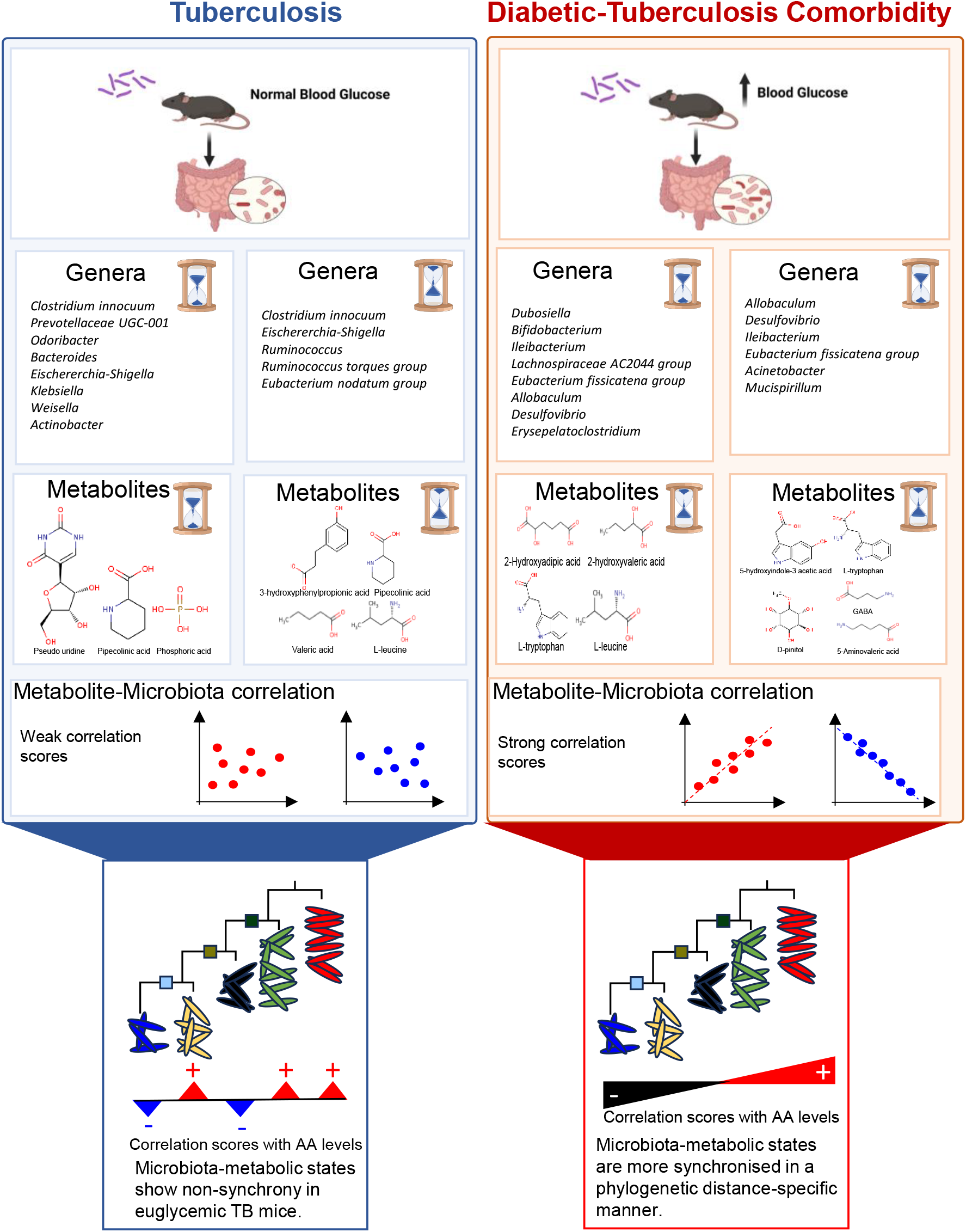
Summary figure. The gut microbiome and metabolome of DM-TB comorbid are condition-specific and different from the TB group.

## Discussion

Diabetes cases are increasing significantly and are becoming a global concern. Countries with higher human development index are reported to show increased DM incidence rates (15,16). DM patients often show systemic metabolic changes in the body, higher levels of advanced glycation end products and chronic inflammation which increases susceptibility to infectious diseases like TB. DM subjects also produce lower IFN-γ and TNF-α levels, critical for controlling Mtb growth inside macrophages. The DMTB comorbid patients reported higher proinflammatory IL-1β, IL-6, and IL-18 cytokines and the macrophages, when exposed to high blood sugar levels, compromised their Mtb phagocytic ability. All these perturbed immune systems in DM result in poor prognosis and may affect Mtb dissemination, causing more severe miliary TB and contributing to drug resistance development (17).

The microbiota plays a crucial role in immune system development. However, factors such as prolonged or unnecessary antibiotic use, dietary changes, or systemic constraints caused by conditions like diabetes, along with genetic factors such as mutations in NOD2 (which prevents gut inflammation) and ATG16L1 (which aids in exocytosis in Paneth cells), can lead to dysbiosis and negatively affect immune function. Dysbiosis in diseased states like inflammatory bowel disease leads to a decrease in alpha diversity, for example, lower *Ruminococcaceae* and *Lactobacillus* abundance and higher *Proteobacteria*. The intricate microbiota-immune system crosstalk in a diseased state can drive changes in Th17, Th2, and Th1 cell counts and decrease Treg and IgA levels, further exacerbating the pathogenesis (18). Metabolites derived from microbes may act as conduits for the crosstalk between microbiota and the immune system. In diabetes, a leaky gut is commonly reported and contributes to generating autoantibodies, resulting in a higher systemic inflammation (19,20). Gut dysbiosis in DM or TB patients has already been reported. However, studies on DM-TB comorbid conditions are limited and need a better understanding (12,21,22).

In the present study, we focused on the metabolome and microbiome landscape of DMTB comorbidity and TB in murine models. Several genera of gut microbiota showed perturbations in their relative proportions in the DMTB group (Figure 7). A decreased alpha diversity in TB and DM patients, compared to healthy subjects, was reported in the literature (23–26). Few studies also reported non-significant changes in alpha diversity in DM patients, indicating the involvement of other factors, possibly due to multiple experimental factors (27,28). In this study, an increase in alpha diversity was observed at 3 weeks in the DM group. In DM patients and obese subjects, a lower abundance of *Akkermansia* was reported, and similar findings were observed in this mice study (29). Corroborating with previous literature, we consistently observed altered beta diversity in the DM group (27). At the genus level, *Allobaculum*, was low in the DM group, which has an intestinal protective role and a reduced abundance of it was reported in the insulin resistance cases (29). An increased abundance of *Allobaculum* was observed in DMTB compared to TB, which seems to be mediated by DM. Higher *Allobaculum* abundance in mice with diabetic nephropathy was reported earlier (30). Genera such as *Desulfovibrio* and *Acinetobacter* are also reported positively associated with hyperglycemia observed in DM or gestational DM (31). *Desulfovibrio* is a sulphate-reducing genus that produces hydrogen sulfide, which contributes to an increase in inflammation (32,33). In the DMTB group, a higher abundance of *Desulfovibrio* was observed, suggesting the change is majorly contributed by diabetes. We observed a higher abundance of *Acinetobacter* in the diseased states (DMTB/TB; DMTB/HC at 8 wpi and TB/HC, 3 wpi), corroborating earlier reports that showed diabetes as a risk factor for *Acinetobacter baumannii* infection (34,35).

Diabetes may impact the microbiome-associated metabolomic landscape. For example, pipecolinic acid, a lysine catabolism product that gut microbes produce, was lower in the DMTB group (36). A negative correlation of serum pipecolinic acid levels with high blood glucose levels in diabetic retinopathy patients and coronary artery disease-diabetic comorbid subjects has already been reported earlier (37,38). At 8 wpi low pipecolinic acid and *Ruminococcus* abundance were observed in the DMTB group compared to the TB group, highlighting their correlation. L-tryptophan was higher in the DMTB group, which gets converted to other secondary metabolites like tryptamine and indoles like indole-3-aldehyde, indole-3-acetic-acid and indole-3-propionic acid by the gut microbiota (39,40). The altered microbial diversity observed in the DMTB group might contribute to higher tryptophan levels. Dysbiosis in DM leads to decreased levels of short-chain fatty acids (SCFA)-producing gut bacteria and reduced SCFAs (viz., propionic acid and butyric acid) (41). SCFAs provide protective functions like promoting anti-inflammatory effects and immune responses. The metabolite-microbiota correlation analysis showed a link between SCFA and valeric acid and the microbes that degrade lactate and produce SCFAs as its by-products, like *Coriobacteriaceae UGC-002. Coriobacteriaceae UGC-002*, belonging to the *Actinobacteria* phylum, is an SCFA-producing bacteria (Figure 7). *Coriobacteriaceae* is reported to be negatively correlated to hyperglycemia observed in diabetes, while it is positively correlated in TB (42–44). A negative correlation between valeric acid and *Coriobacteriaceae UGC-002* was observed between the DMTB and TB group at 3 wpi. The reduced valeric acid levels in TB with an increased *Coriobacteriaceae* level indicate its impact on DMTB comorbidity. In a recent report, the administration of probiotics in T2DM patients increased valeric acid levels, and a similar correlation between valeric acid and similar probiotic bacteria was observed in this study (45).

Microbiota also demonstrates metabolic state synchrony by coordinating the gut microbiome and metabolism with circadian rhythm (46,47). We observed correlating amino acid levels and microbiota in a phylogenetic distance-specific manner, indicating possible synchrony in microbial metabolism for selected genera. Previous reports indicated that gut tryptophan metabolism affects the glycemic state through peripheral serotonin, impacting host homeostasis (48). Also, the involvement of intestinal microbiota in amino acid utilization for SCFA production and the impact of T2DM on it is well-known (49). Our findings highlight higher synchrony in amino acid metabolism in the DMTB group and shed light on phylogenetic clades that may have a role in SCFA production and intestine wall protection (Figure 7).

There are a few limitations in this study, including the use of only male mice and the inherent differences in the DM murine model and its translational potential in patients. Further expansion, including female mice and understanding the effect of TB drugs on the studied aspects, will provide more meaningful information, which is ongoing in our laboratory.

## Conclusion

This study highlights distinct gut microbiota dysbiosis and metabolic alterations in the DMTB comorbid condition compared to TB controls. Several microbiota signatures and associated metabolites were identified as potential biomarkers for DMTB, offering promising avenues for the development of targeted therapeutic interventions, including prebiotic and probiotic supplementation.

## Data Availability Statement

The metagenomic raw data files have been deposited to NCBI Sequence Read Archive (SRA) and can be accessed with the BioProject accession PRJNA1253975.

## Contributions

SJ and RKN conceptualized the study. SJ performed animal handling, faecal material collection, sample processing, and data acquisition using mass spectrometry and gDNA isolation. NB and SJ analysed the data, prepared figures, and prepared the first draft of the manuscript.

## Acknowledgements

SJ acknowledges a Junior Research Fellowship from UGC. NB is financially supported by a grant from the Department of Biotechnology (DBT), GoI (Grant ID: National Network Project of National Institute of Immunology, New Delhi-[40267]). SC is SPM fellow supported by CSIR. RKN acknowledges core support from the ICGEB New Delhi Component. Ms. Riya Ahmed and Mr. Anil Behera from the Translational Health Group, ICGEB, New Delhi component, are acknowledged for providing technical help during the study. All authors acknowledge Tuberculosis Aerosol Challenge Facility (TACF) BSL-III of ICGEB, New Delhi component for providing lab space to culture Mtb and perform animal infection studies.

