## supplementary file for "Diabetes mellitus and tuberculosis comorbidity induce unique microbial gut dysbiosis and associated metabolome"

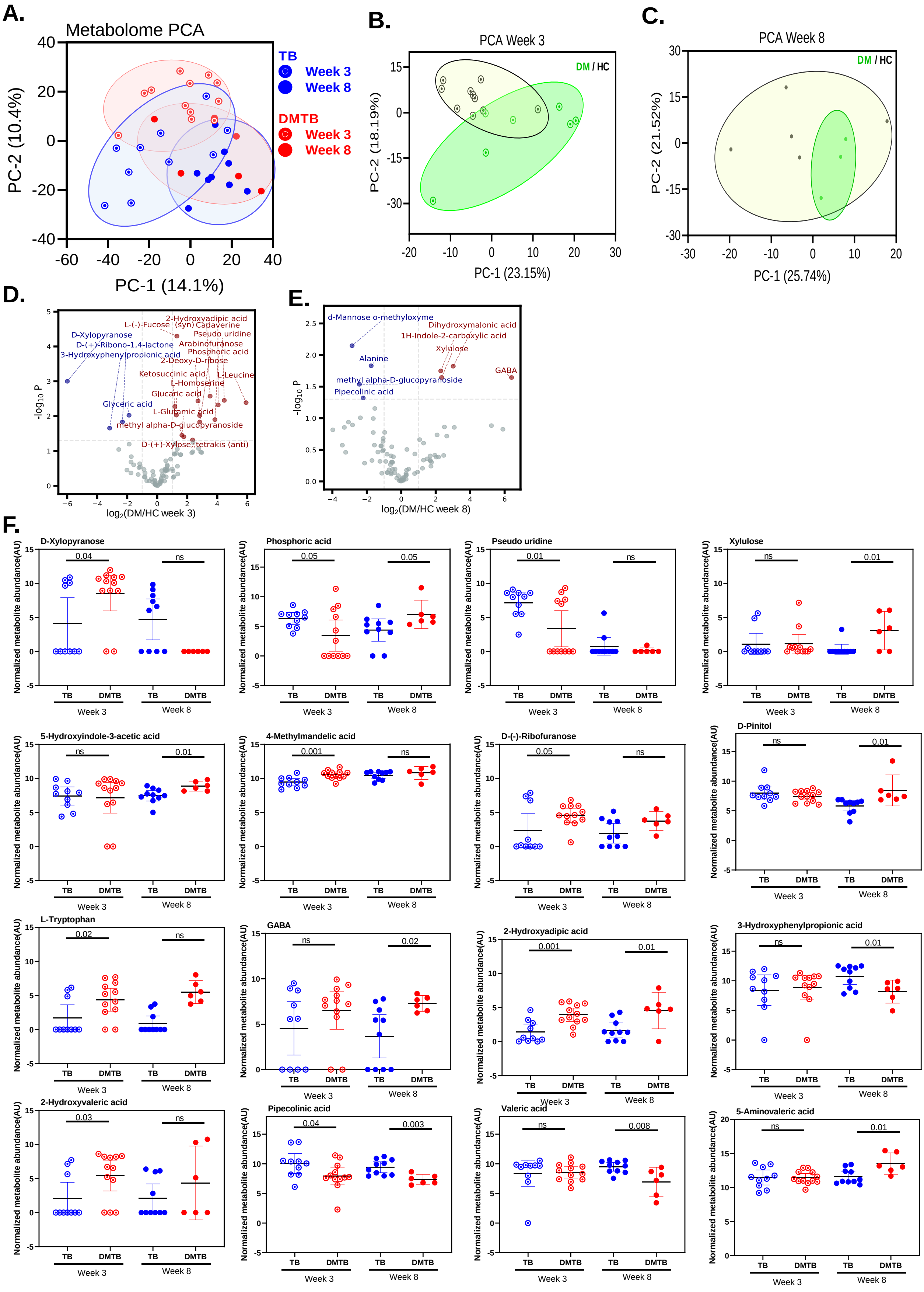


**Figure S1: Fecal Metabolites are impacted by diabetes in C57Bl/6 mice.** The metabolome matrix was log-transformed and subjected to PCA analysis and differential metabolite analysis using a student's t-test. A: PCA plot of DMTB, TB at Week 3 and Week 8 post infection. B: PCA plot of DM and HC at Week 3. C: PCA plot of DM and HC at Week 8. D: Differential abundance analysis of DM vs HC at week 3. E: Differential metabolite analysis of DM vs HC at week 8. F: Relative abundance of metabolites between DMTB and TB at week 3 and week 8 post infection.

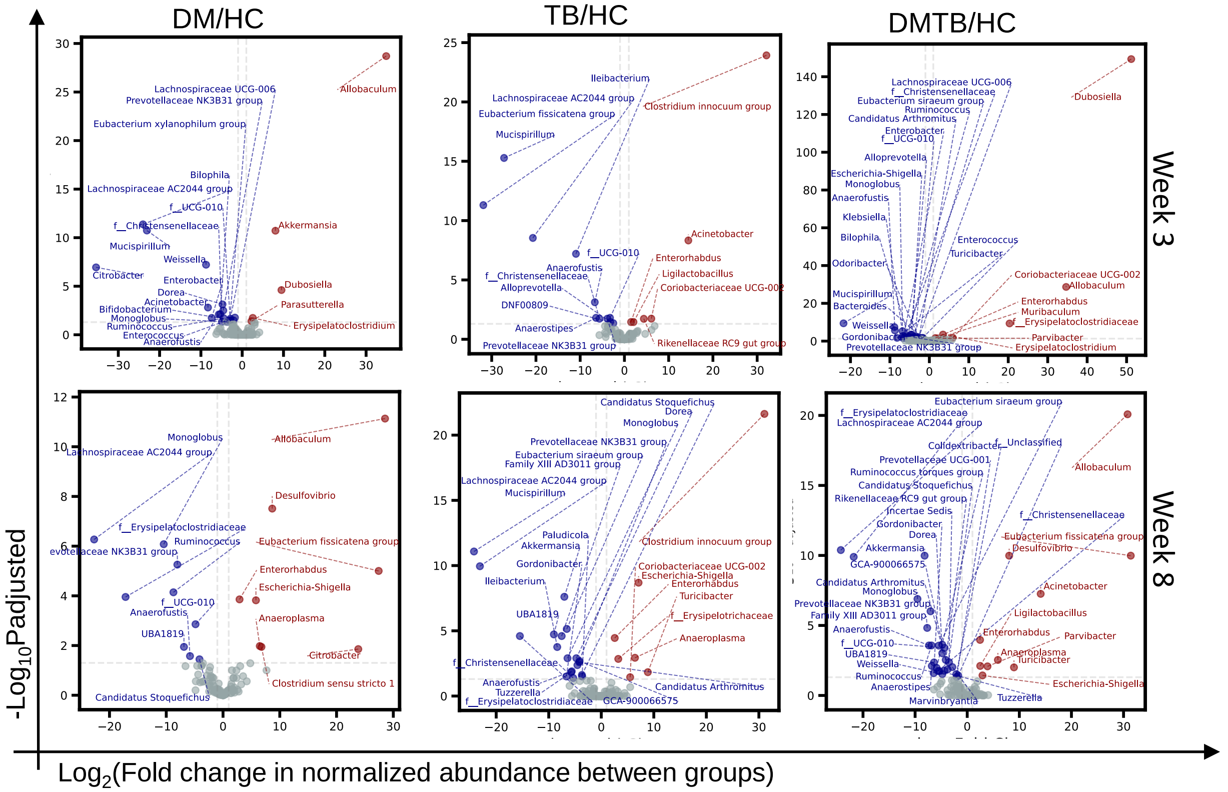


**Figure S2: Fecal microbiome changes are impacted by diabetes after *Mtb* infection.** Differentially abundant genera (Padj<0.05, log2FC>±1) in TB, DM and DMTB.

**A.**

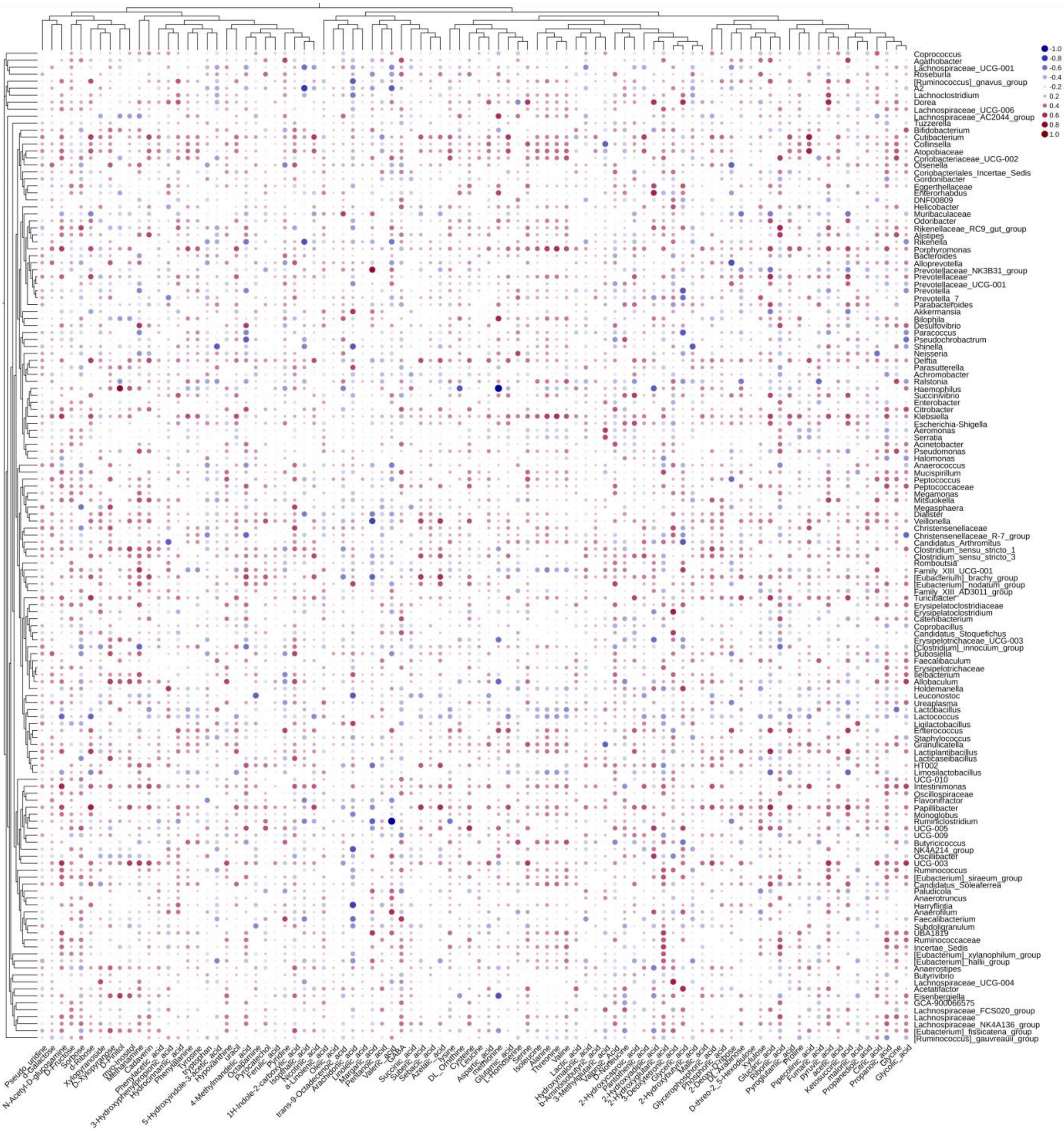


Figure S3: Filtered correlation between metabolites and microbiota. Correlation in TB at Week 3 (A) and Week 8 (B). Correlation in DMTB at week 3 (C) and at week 8 (D).

Figure S3 continued.

**B.**

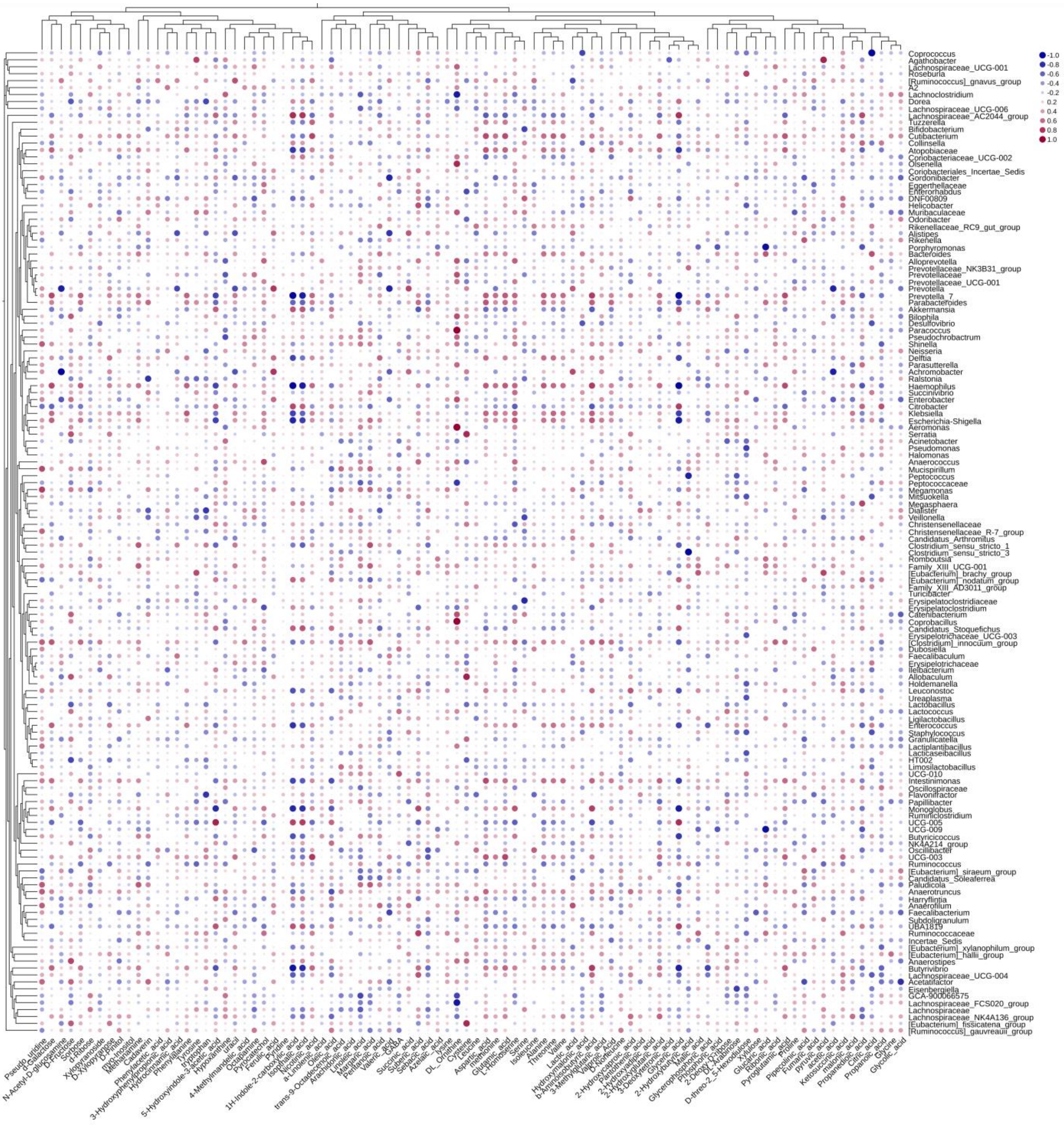

Figure S3 continued.

C.

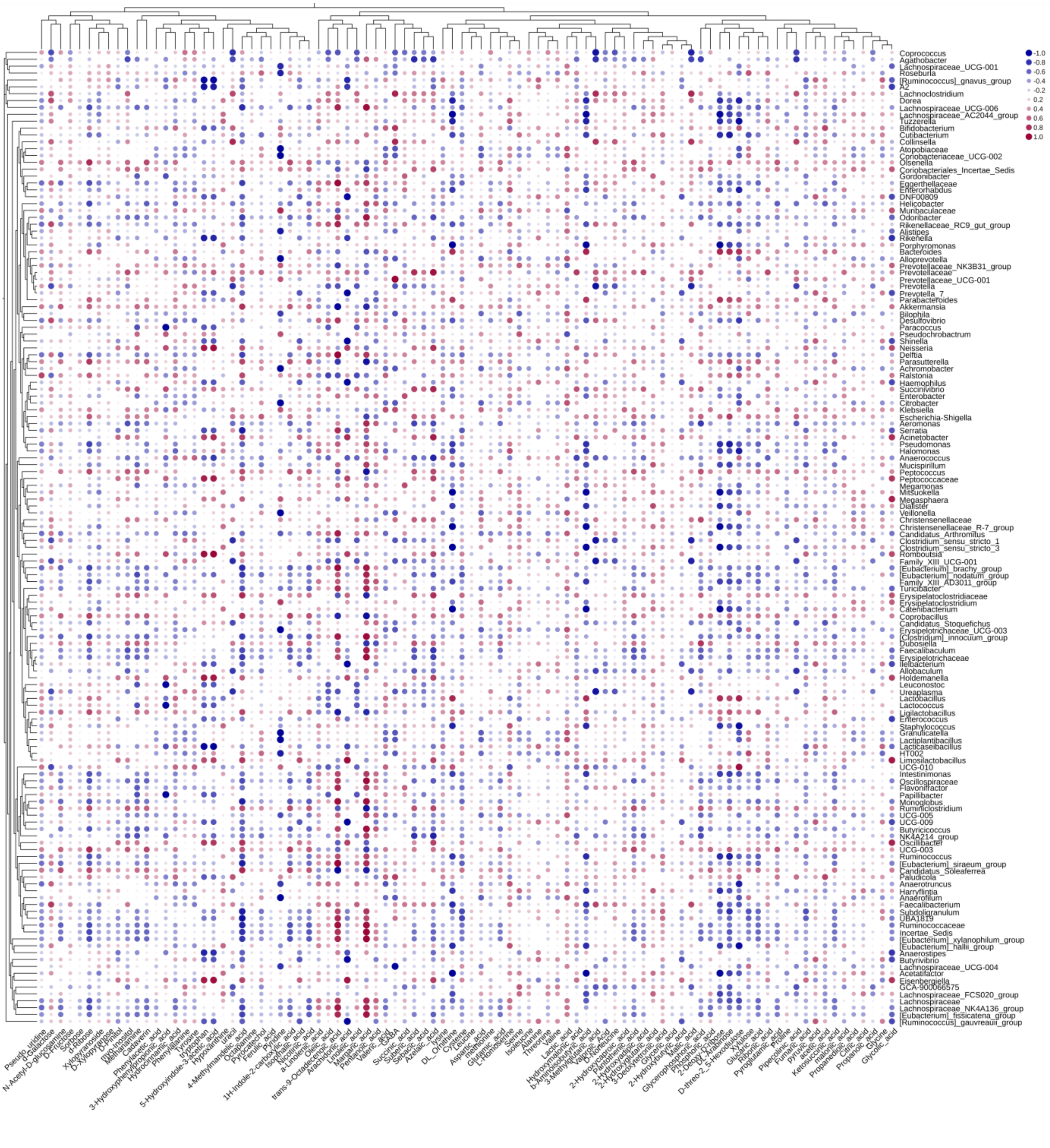

Figure S3 continued.

**D.**

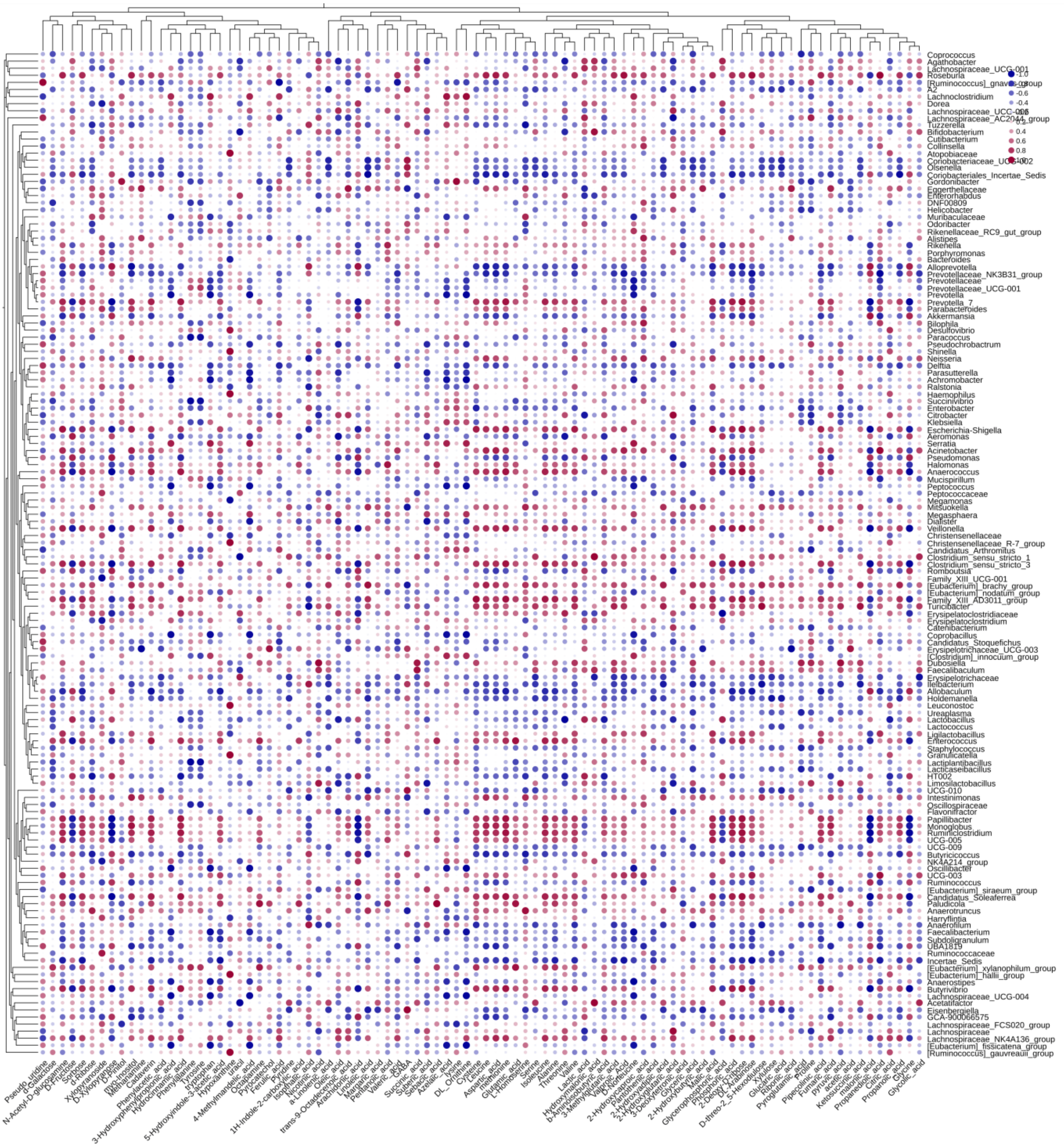
